# A promising QTL *QSns.sau-MC-3D.1* likely superior to *WAPO1* for wheat spikelet number per spike shows no adverse effects on yield-related traits

**DOI:** 10.1101/2023.02.17.528911

**Authors:** Jieguang Zhou, Wei Li, Yaoyao Yang, Xinlin Xie, Jiajun Liu, Yanling Liu, Huaping Tang, Mei Deng, Qiang Xu, Qiantao Jiang, Guoyue Chen, Pengfei Qi, Yunfeng Jiang, Guangdeng Chen, Yuanjiang He, Yong Ren, Liwei Tang, Lulu Gou, Youliang Zheng, Yuming Wei, Jian Ma

**Affiliations:** State Key Laboratory of Crop Gene Exploration and Utilization in Southwest China, Sichuan Agricultural University, Chengdu, China; Triticeae Research Institute, Sichuan Agricultural University, Chengdu, China; College of Resources, Sichuan Agricultural University, Chengdu, China; Mianyang Academy of Agricultural Science/Crop Characteristic Resources Creation and Utilization Key Laboratory of Sichuan Providence, Mianyang, China; Panzhihua Academy of Agricultural and Forestry Sciences, Panzhihua, China; College of Agronomy, Sichuan Agricultural University, Chengdu, China

## Abstract

Spikelet number per spike (SNS) is one of the crucial factors determining wheat yield. Thus, improving our understanding of the genes that regulate SNS could help develop higher-yielding wheat varieties. A genetic linkage map constructed using the GenoBaits Wheat 16K Panel and the 660K SNP array contained 5991 polymorphic SNP markers spanning 2813.26 cM. A total of twelve QTL for SNS were detected in the recombinant inbred line (RIL) population *msf* × Chuannong 16 (MC), and two of them, i.e., *QSns.sau-MC-3D.1* and *QSns.sau-MC-7A*, were stably expressed. *QSns.sau-MC-3D.1* had high LOD values ranging from 4.99 to 11.06 and explained 9.71-16.75% of the phenotypic variation. Comparison of *QSns.sau-MC-3D.1* with previously reported SNS QTL suggested that it is likely a novel one. A kompetitive allele-specific PCR (KASP) marker, KASP-10, tightly linked to *QSns.sau-MC-3D.1* was developed to successfully validate its effect in three segregated populations and a natural population. Genetic analysis indicated that *WHEAT ORTHOLOG OFAPO1* (*WAPO1*) was a candidate gene for *QSns.sau-MC-7A*. The combined additive effect of *QSns.sau-MC-3D.1* and *WAP01* had a great additive effect increasing SNS by 7.10%. In addition, our results suggested that SNS is not affected by 1BL/1RS translocations in the MC RIL population. Correlation analysis between two major QTL and other agronomic traits showed that *QSns.sau-MC-3D.1* was likely independent of these agronomic traits. However, the H2 haplotype of *WAPO1* may affect effective tiller number and plant height. This indicated that the breeding potential of *QSns.sau-MC-3D.1* is better than that of *WAPO1*. The geographical distribution of *QSns.nsau-MC-3D.1* showed that *QSns.sau-MC-3D.1* positive allele frequency was dominant in most wheat-producing regions of China and it has been positively selected among modern cultivars released in China since the 1940s. Two genes, *TraesCS3D03G0222600* and *TraesCS3D03G0216800*, associated with SNS development were predicted in the physical interval of *QSns.sau-MC-3D.1*. qRT-PCR results of the two genes showed that only the expression level of *TraesCS3D03G0216800* was significantly different between msf and CN16. These results enrich our understanding of the genetic basis of wheat SNS and will be useful for fine mapping and cloning of genes underlying *QSns.sau-MC-3D.1*, and provide a basis for marker-assisted selection breeding.

**Author summary:** In this study, we identified two major QTL (*QSns.sau-MC-3D.1* and *QSns.sau-MC-7A*) in a RIL population. *WAPO1* was demonstrated to be the candidate gene for *QSns.sau-MC-7A. QSns.sau-MC-3D.1* was a novel and stably expressed QTL, and further confirmed in different genetic backgrounds. Our results further demonstrate that *QSns.sau-MC-3D.1* has better breeding potential because of its no adverse effect on other agronomic traits than *WAPO1*, and it has been positively selected during Chinese breeding programs since the 1940s. Taken together, the identification of *QSns.sau-MC-3D.1* offers a promising resource to further increase wheat yields.

## Introduction

Bread wheat (AABBDD, *Triticum aestivum* L.) is one of the most important food crops in the world [1]. The increasing population and frequent natural disasters [2] lead to the world confronting a huge food gap, and high yield has always been the eternal theme of wheat breeding. Kernels per spike (KNS), thousand kernel weight (TKW), and spikes per unit area are the three components of yield [3,4]. Spikelet number per spike (SNS) is closely related to KNS [5], and breeders can usually improve wheat yield by increasing SNS. Thus, it is essential to understand the genetic pattern of SNS for optimizing wheat spike structure and cultivating new high-yielding wheat varieties.

To date, quantitative trait loci (QTL) of SNS have been detected on all 21 chromosomes of wheat using bi-parental populations [6]. For example, Zhai et al. [7] used the RIL population to detect a major QTL on chromosome 1B controlling SNS, which explained 30.75% of the phenotypic variance (PVE). *QSns.sau-2D* on chromosome 2D significantly increased SNS by up to 14.72% [8]. Mo et al. [9] identified two major and novel SNS-related QTL, *QSns.sau-AM-2B.2* and *QSns.sau-AM-3B.2*, using a tetraploid RIL population. The SNP marker Kukri_c8913_655, which is tightly linked to SNS, was identified on chromosome 3D [7]. Furthermore, some genes related to SNS have been reported, such as *trs1*/*WFZP-A* [10], *VRN-A3/FT-A1* [11], *Q* [12], *TaTB1-4A* [13], *PPD-A1* [14], *TaCol-B5* [15], and *WHEAT ORTHOLOG OFAPO1* (*WAPO1*) [16-19]. Although many QTL/genes associated with SNS have been reported in wheat, major, stably expressed and confirmed QTL in multiple environments and genetic backgrounds and high-efficiency molecular markers is still limited.

Single nucleotide polymorphisms (SNPs) are the most abundant and important type of nucleic acid variation [20]. To date, multiple SNP arrays have been developed in wheat, such as the 9K, 16K, 55K, 90K, 660K, and 820K high-density SNP chips. The Wheat 16K array was developed using an improved genotyping by target sequencing (GBTS) system with capture-in-solution (liquid chip) technology [21]. The 16K SNP was identified based on resequencing data from 20 accessions, genotyping data of 1,520 germplasms collected from multiple platforms, and publicly released resequencing and exon capture data. These SNP datasets were developed and optimized using GenoBait technology to eventually produce 14,868 multiple SNP (mSNP) segments (including 37,669 SNP markers) [22].

In this study, we report a genetic map of bread wheat constructed based on the Wheat 16K SNP array and the Wheat 660K SNP array. Using the constructed genetic map, QTL for SNS were identified. Major and novel QTL were validated in four populations with different genetic backgrounds *via* kompetitive allele-specific PCR (KASP) markers. Furthermore, the genetic effects and geographical distribution of the major QTL were also analyzed to clarify their application potential in breeding and to provide a theoretical basis for genetic improvement of wheat yield.

## Results

### SNP markers and genetic linkage map

Of the 37,671 SNPs, 5,991 (∼15.90%) with MAF ≥0.3 and showing polymorphisms between parents in the MC mapping population were retained for further analysis. These 5,991 SNP markers were divided into 1,198 bins using the ‘BIN’ function in QTL IciMapping V4.1 and markers with the lowest ‘missing rate’ in each bin (bin markers) were selected and used to construct the genetic map (Table 1). The resultant linkage map consisted of 1,150 bin markers classified into 26 linkage groups (Table 1). Among them, chromosomes 3D, 4A, 5A, 5D, and 7B each had two linkage groups, and only one was constructed for each of the remaining chromosomes (Table 1). Chromosome arm 1BS was not covered by any marker mainly due to the 1BL/1RS translocation on chromosome 1B (Fig 1). The total length of the 26 linkage groups was 2,813.26 cM, with an average spacing of 2.45 cM (Table 1). The A, B, and D genomes included 479 (∼41.65%), 473 (∼41.13%), and 198 (∼17.22%) markers covering lengths of 1,047.05, 925.92, and 840.28 cM with average marker intervals of 2.19, 1.96, and 4.24 cM, respectively (Table 1). The lowest marker coverage was detected for the D genome, especially for chromosomes 4D and 6D (Table 1).

**Table 1.** Details of markers in the constructed genetic map.

| Chromosome | Group | Number of bin marker | Number of mapped markers | Length (cM) | cM per bin marker |
| --- | --- | --- | --- | --- | --- |
| 1A | 1 | 78 | 546 | 131.19 | 1.68 |
| 1B | 1 | 47 | 557 | 93.11 | 1.98 |
| 1D | 1 | 37 | 150 | 119.24 | 3.22 |
| 2A | 1 | 77 | 297 | 148.17 | 1.92 |
| 2B | 1 | 77 | 414 | 209.96 | 2.73 |
| 2D | 1 | 31 | 132 | 104.03 | 3.36 |
| 3A | 1 | 84 | 439 | 185.41 | 2.21 |
| 3B | 1 | 74 | 427 | 145.19 | 1.96 |
| 3D | 2 | 43 | 101 | 250.45 | 5.82 |
| 4A | 2 | 57 | 318 | 147.74 | 2.59 |
| 4B | 1 | 46 | 169 | 122.34 | 2.66 |
| 4D | 1 | 23 | 69 | 54.55 | 2.37 |
| 5A | 2 | 84 | 362 | 161.70 | 1.92 |
| 5B | 1 | 64 | 258 | 137.45 | 2.15 |
| 5D | 2 | 29 | 78 | 153.05 | 5.28 |
| 6A | 1 | 48 | 327 | 141.65 | 2.95 |
| 6B | 1 | 77 | 751 | 84.71 | 1.10 |
| 6D | 1 | 7 | 41 | 60.91 | 8.70 |
| 7A | 1 | 51 | 171 | 131.20 | 2.57 |
| 7B | 2 | 88 | 313 | 133.16 | 1.51 |
| 7D | 1 | 28 | 71 | 98.05 | 3.50 |
| A genome | 9 | 479 | 2460 | 1047.05 | 2.19 |
| B genome | 8 | 473 | 2889 | 925.92 | 1.96 |
| D genome | 9 | 198 | 642 | 840.28 | 4.24 |
| Total | 26 | 1150 | 5991 | 2813.25 | 2.45 |

**Fig 1.**
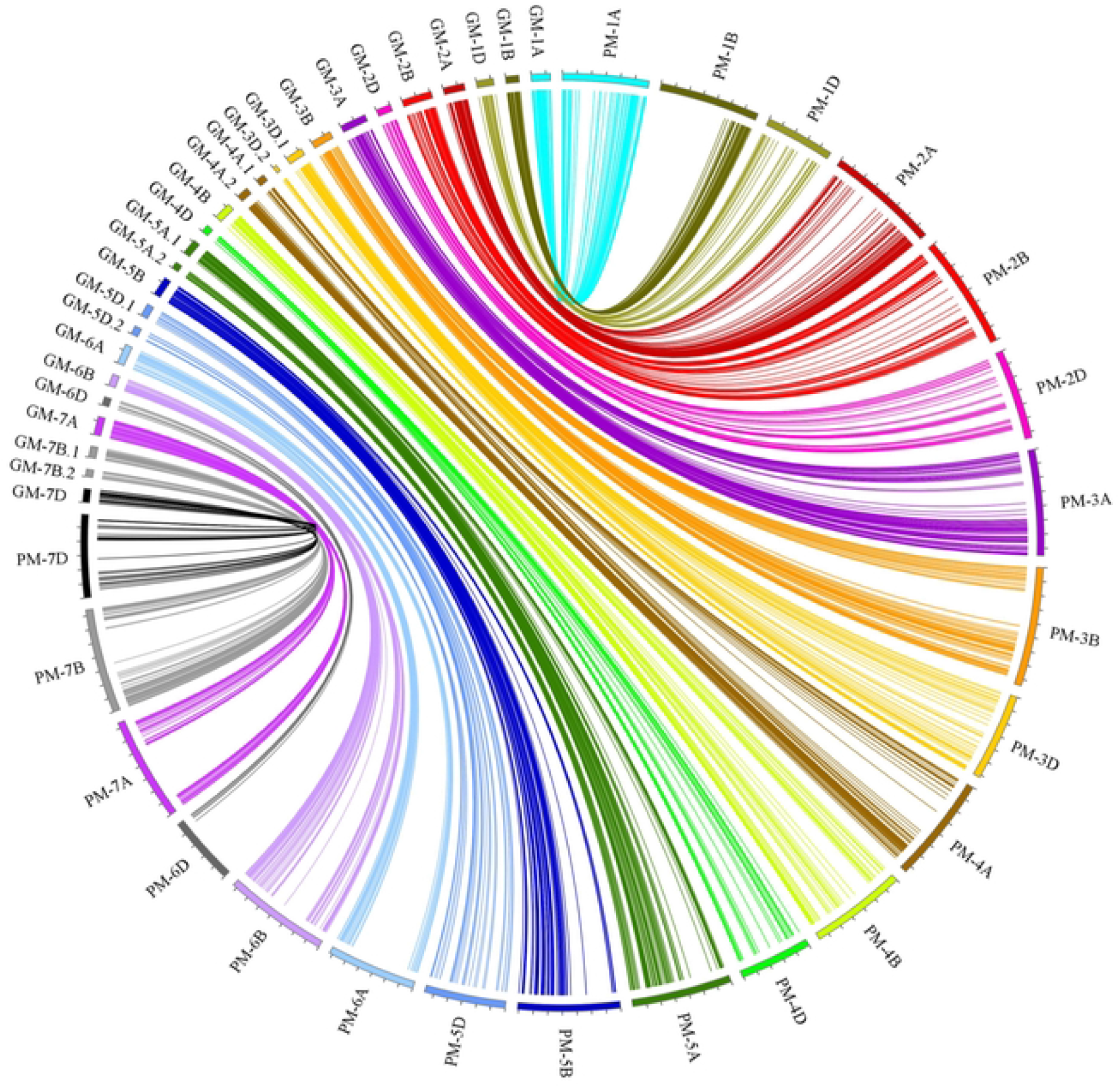
The syntenic relationships between the genetic and physical maps of bin markers. GM-1A to GM-7D represented the 26 chromosomal genetic maps used in this study; PM-1A to PM-7D represented the 21 chromosomal physical maps of wheat.

### Comparison of genetic and physical maps

The sequences of the 5,991 mapped markers were blasted against CS V2.1 genome to obtain their physical positions (S1 Table). Among them, 5980 markers (99.82%) showed coincident physical and genetic positions (S2 Table). The genetic positions of the 1,150 bin markers were compared with their physical positions in the CS V2.1 genome, and 1,015 (∼88.26%) markers showed good concordance (Fig 1 and S3 Table).

### Phenotypic variation and ANOVA in all environments

*msf* had a higher value of SNS than CN16 (*P* < 0.01) in five environments (Table 2). The SNS of the MC RIL population ranged between 14.00 and 29.00 and was normally and continuously distributed (S1 Fig and Table 2), indicating polygenic control. The estimated *H*^2^ of SNS was 0.74, indicating that SNS was significantly affected by genetic factors (Table 2). ANOVA showed a significant effect of G (Genotype), E (Environment), and G × E interaction on SNS (*P* < 0.001; S4 Table). However, Block/E did not differ significantly (*P* > 0.05) on SNS (S4 Table), suggesting that two planting replicates within a single environment were reliable and meaningful.

**Table 2.** Phenotypic variation of spikelet number per spike (SNS) for the mapping population *msf* × CN16 and parental lines in five environments and BLUP.

| Env. | Parents |  | *msf* × CN16 |  |  |  |
| --- | --- | --- | --- | --- | --- | --- |
| | *msf* | CN16 | Min–Max | Mean | SD | $H^2$ |
| 2021WJ | 27.60** | 20.00 | 19.00–26.75 | 23.08 | 1.64 |  |
| 2021CZ | 25.50** | 20.44 | 19.00–25.88 | 22.78 | 1.26 |  |
| 2021YA | 27.67** | 22.00 | 18.00–29.00 | 23.45 | 1.97 |  |
| 2022WJ | 24.00** | 18.13 | 17.14–26.00 | 21.06 | 1.58 |  |
| 2022CZ | 24.17** | 19.00 | 14.00–28.00 | 21.35 | 2.09 |  |
| BLUP | 25.12** | 20.30 | 19.89–24.42 | 21.85 | 0.96 | 0.74 |

Env., Environment; \*\*Significance level at $P < 0.01$ ; SD, standard deviation; $H^2$ , broad-sense heritability; WJ, Wenjiang; CZ, Chongzhou; YA, Ya'an; BLUP, best linear unbiased prediction.

### Correlation analyses between SNS and other agronomic traits

SNS showed a significant positive correlation (*P* < 0.01) in all five environments and BLUP dataset (Fig 2), with coefficients ranging from 0.32 and 0.79 (Fig 2). The BLUP dataset of SNS and five other agronomic traits were employed to evaluate their Pearson’s correlations. There was a significant correlation between SNS and SL, AD, and TKW (Table 3). Furthermore, there was no significant correlation between SNS and ETN, and PH (Table 3).

**Table 3.** Correlations between spikelet number per spike (SNS) and other agronomic traits in the mapping population *msf* × CN16 population.

| Trait | Spikelet number per spike (SNS) |
| --- | --- |
| Effective tiller number (ETN) | -0.036 |
| Plant height (PH) | 0.086 |
| Spike length (SL) | 0.35** |
| Anthesis date (AD) | 0.40** |
| Thousand kernel weight (TKW) | -0.14* |

\*Significance level at $P < 0.05$ ; \*\*significance level at $P < 0.01$ .

**Fig 2.**
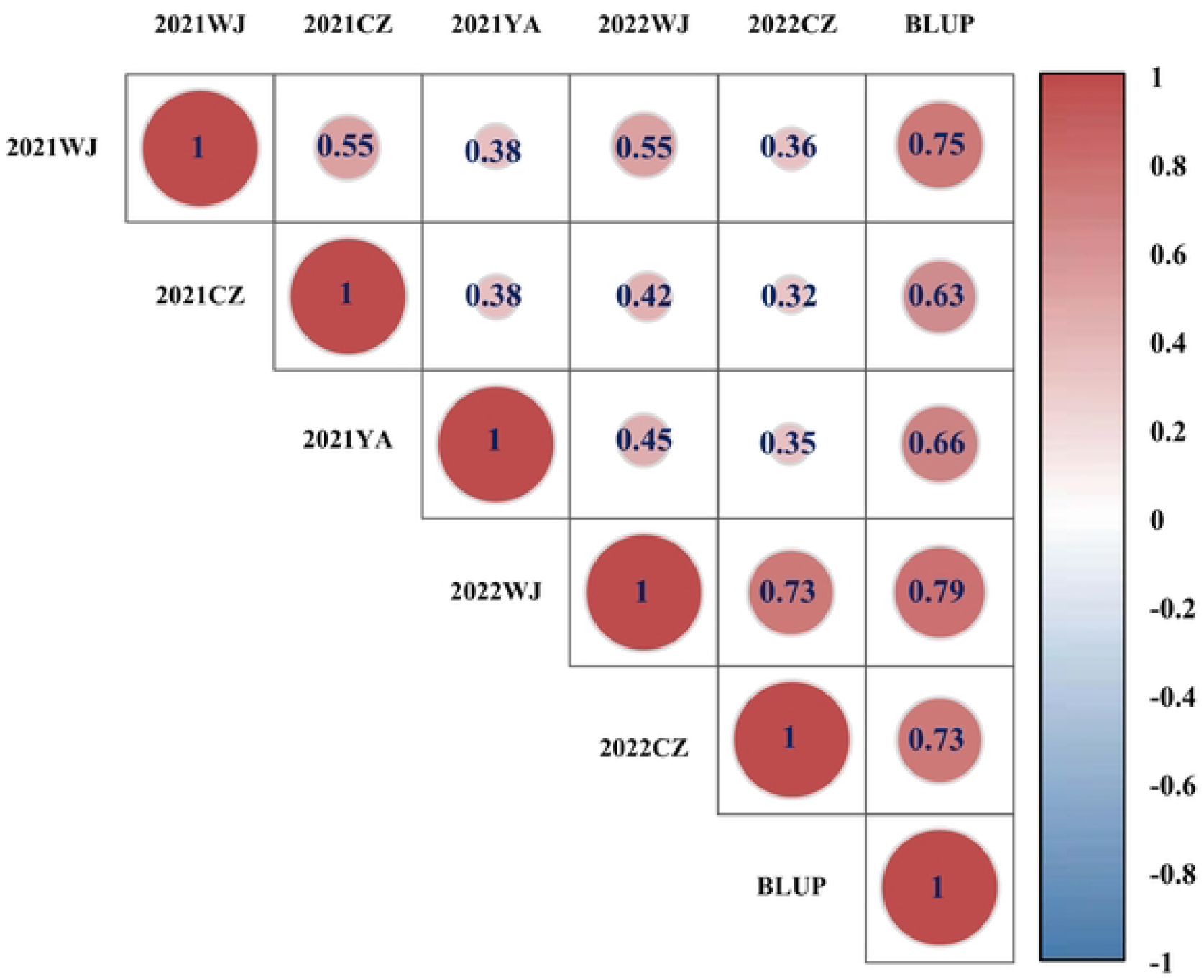
The correlation coefficients of spikelet number per spike (SNS) in multiple environments. The blue-colored ‘correlation coefficient’ represents a significant level at *P*<0.01.

### Identification of QTL for SNS

Twelve QTL for SNS were identified, and they were located on chromosomes 1B, 2A, 3D (2), 4A, 5A, 5B (2), 6A, 6B, 7A, and 7B, with LOD scores ranging between 2.52 and 16.66 (Table 4). Among them, *QSns.sau-MC-3D.1* and *QSns.sau-MC-7A* were identified in three and five environments as well as using the BLUP dataset (Table 4), respectively. Therefore, these two QTL were considered as the major and stable QTL for SNS. The remaining eight QTL were detected in a single or two environments explaining between 3.21% and 9.61% of the PVE and they were accordingly designated as minor QTL (Table 4). The positive alleles of *QSns.sau-MC-3D.1* and *QSns.sau-MC-7A* were both derived from the parent *msf*.

**Table 4.** Quantitative trait loci for spikelet number per spike (SNS) identified in the mapping population *msf* × CN16 evaluated in five environments and BLUP.

| QTL | Env. | Interval (cM) | Flanking marker | LOD | PVE (%) | Add | Physical position (Mb) |
| --- | --- | --- | --- | --- | --- | --- | --- |
| *QSns.sau-MC-1B* | 2021CZ | 84.5-93 | 1B_679421143-1B_687066854 | 2.65 | 4.26 | -0.27 | 688.30-696.94 |
| *QSns.sau-MC-2A* | BLUP | 53.5-64.5 | 2A_144065493-2A_165841608 | 2.52 | 3.21 | 0.18 | 148.72-170.38 |
| *QSns.sau-MC-3D.1* | 2021YA | 81.5-89.5 | KASP-10-3D_64273412 | 7.2 | 16.75 | 0.82 | 53.61-64.40 |
|  | 2022WJ | 83.5-88.5 | KASP-10-3D_64273412 | 9.71 | 14.82 | 0.62 |  |
|  | 2022CZ | 81.5-88.5 | KASP-10-3D_64273412 | 4.99 | 9.71 | 0.63 |  |
|  | BLUP | 83.5-88.5 | KASP-10-3D_64273412 | 11.06 | 16.34 | 0.4 |  |
| *QSns.sau-MC-3D.2* | 2021WJ | 96.5-100.5 | 3D_122396589-3D_138793245 | 3.05 | 5.32 | 0.38 | 122.97-139.28 |
| *QSns.sau-MC-4A* | 2022WJ | 29.5-35.5 | 4A_639942192-4A_679248111 | 6.31 | 9.61 | 0.5 | 639.38-681.19 |
|  | 2022CZ | 29.5-35.5 | 4A_639942192-4A_679248111 | 4.41 | 8.47 | 0.60 |  |
|  | BLUP | 28.5-35.5 | 4A_639942192-4A_679248111 | 2.95 | 3.85 | 0.19 |  |
| *QSns.sau-MC-5A* | 2022WJ | 0-13.5 | 5A_9510718-5A_27776458 | 2.57 | 4.42 | 0.34 | 11.56-29.72 |
| *QSns.sau-MC-5B.1* | 2021WJ | 9.5-17.5 | 5B_38648213-5B_43372176 | 3.48 | 6.16 | 0.41 | 39.68-230.88 |
|  | BLUP | 10.5-19.5 | 5B_43372176-5B_227699843 | 3.45 | 5.06 | 0.22 |  |
| *QSns.sau-MC-5B.2* | 2021CZ | 24.5-25.5 | 5B_401661494-5B_414921497 | 2.85 | 4.59 | 0.28 | 404.64-417.83 |
| *QSns.sau-MC-6A* | 2022WJ | 0-5.5 | 6A_3774779-6A_5558438 | 3.16 | 4.37 | 0.34 | 4.57-6.86 |
| *QSns.sau-MC-6B* | 2022CZ | 23.5-24.5 | 6B_644045066-6B_647440241 | 2.99 | 5.34 | -0.47 | 652.23-655.69 |
| *QSns.sau-MC-7A* | 2021WJ | 100.5-107.5 | 7A_671413788-7A_672390144 | 7 | 12.85 | 0.59 | 676.01-679.91 |
|  | 2021CZ | 105.5-108.5 | 7A_671413788-7A_672390144 | 12.28 | 22.42 | 0.61 |  |
|  | 2021YA | 107.5-109.5 | 7A_672390144-7A_673311365 | 7.32 | 15.87 | 0.8 |  |
|  | 2022WJ | 100.5-107.5 | 7A_671413788-7A_672390144 | 8.55 | 12.71 | 0.57 |  |
|  | 2022CZ | 101.5-107.5 | 7A_671413788-7A_672390144 | 4.97 | 8.95 | 0.61 |  |
|  | BLUP | 105.5-107.5 | 7A_671413788-7A_672390144 | 16.66 | 24.25 | 0.49 |  |
| ***QSns.sau-MC-7B*** | 2021WJ | 73.5-80 | 7B_582312831-7B_587920157 | 2.77 | 4.79 | 0.36 | 586.93-592.64 |

QTL, quantitative trait loci; Env., environment; BLUP, best linear unbiased prediction; LOD, logarithm of odds; PVE, phenotype variance explained. Add, additive effect (positive values: positive alleles from *msf*, negative values: positive alleles from CN16). Physical position, the flanking marker sequences of QTL were blasted against IWGSC RefSeq V2.1 to get physical positions.

*QSns.sau-MC-3D.1* was located in an 8-cM region between KASP-10 and 3D_64273412. It explained 9.71-16.75% of the PVE (Table 4). The effect of *QSns.sau-MC-3D.1* was highly significant (*P* < 0.01) in five environments and BLUP dataset (Fig 3B). According to flanking marker profiles of *QSns.sau-MC-3D.1*, lines with the homozygous genotype GG from *msf* had significantly higher (*P* < 0.01) SNS than those with the homozygous genotype AA from CN16 and the difference ranged from 2.29 to 6.94% (Fig 3B).

**Fig 3.**
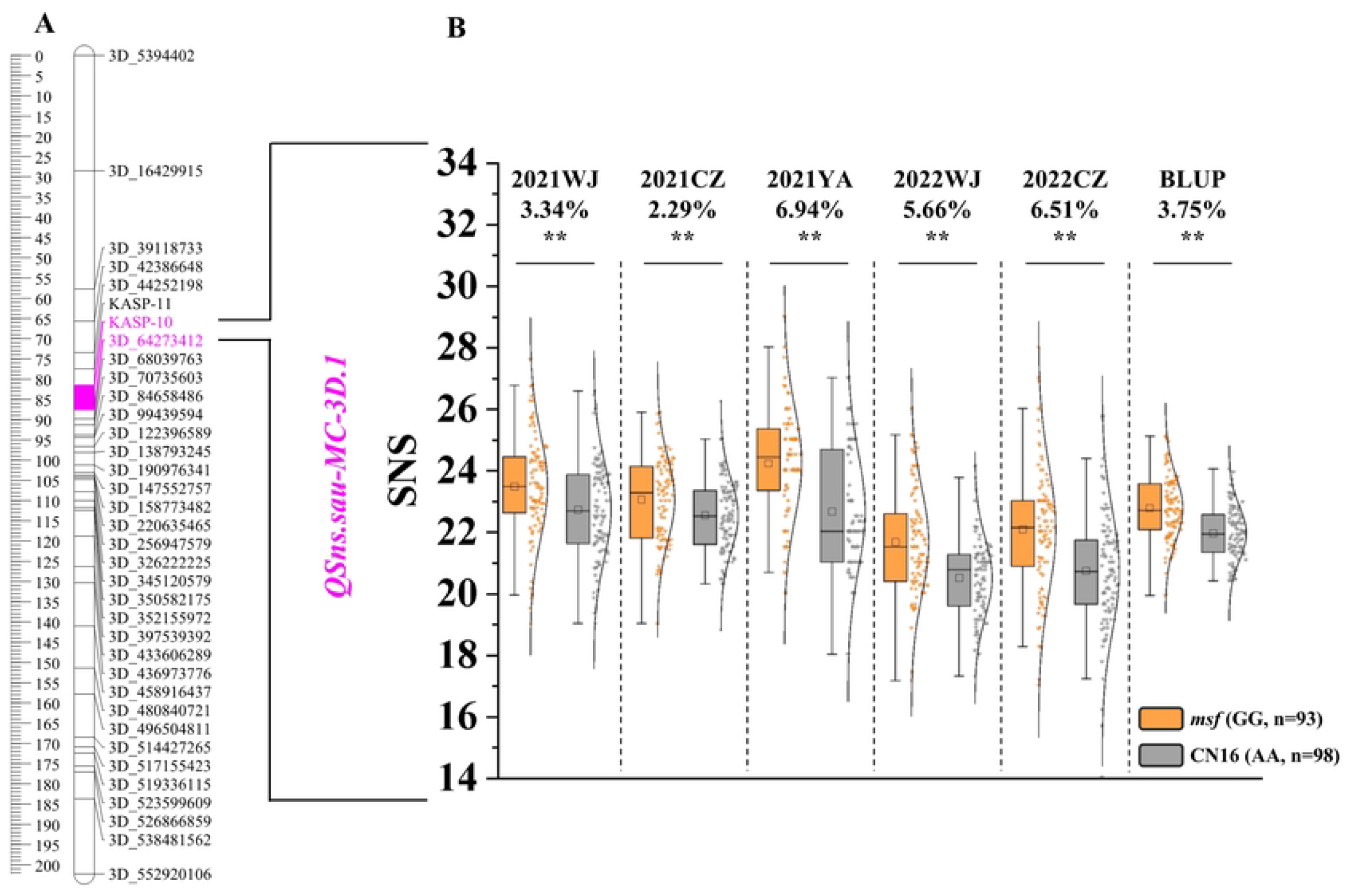
The genetic map of the major *QSns.sau-MC-3D.1* and its effect. **A**, Genetic map of chromosome 3D. The red area is the interval of *QSns.sau-MC-3D.1*. **B**, A box plot that shows the effect of *QSns.sau-MC-3D.1* calculated after grouping the MC RIL population into two categories based on the genotypes of flanking markers. Orange and grey boxes indicate lines with the homozygous genotype from *msf* (GG) and CN16 (AA), respectively. **Significance level at *P* < 0.01, ns indicates no significant difference between the two groups. Differences in SNS between the two groups are labeled below the environment names and BLUP.

*QSns.sau-MC-7A* was stably detected in all environments and located in a 9-cM region between 7A_671413788 and 7A_673311365 (Table 4). It can explain up to 24.25% of the PVE (Table 4). *QSns.sau-MC-7A* was located between 676.00 and 679.91 Mb on CS 7AL by anchoring flanking markers 7A_671413788 and 7A_673311365 (Table 4). Here, it is worth noting that *WAPO1* (*TraesCS7A03G1166400*) is also located in this interval [19]. According to previous studies by Kuzay et al. [19] and Ding et al. [23], *WAPO1* was classified into four haplotypes, including H1 (140^G^+115^deletion^), H2 (140^T^+115^insertion^), H3 (140^G^+115^insertion^), and H4 (140^T^+115^deletion^), based on the types of SNP in its F-box region and a insertion/deletion fragment in the promoter sequence. Hence, we used the previously reported functional marker (K-WAPO1) and Indel marker (WAPO1-ProS) of *WAPO1* for genotyping *msf* and CN16 (S5 Table). Genotyping results showed that *msf* and CN16 belong to H2 and H3, respectively (S2 Fig). This result is consistent with the previous result that H2 is an excellent haplotype that can increase SNS [23], and further suggests that *WAPO1* is likely the causal gene for *QSns.sau-MC-7A*. Furthermore, the MC RIL population was divided into two categories (lines with haplotypes H2 and H3, respectively) based on the genotyping result of K-WAPO1. SNS of the category with H2 had significantly (*P* < 0.01) greater values than that with H3 in each environment and BLUP dataset (S3B Fig).

### Validation of *QSns.sau-MC-3D.1*

The effects of *QSns.sau-MC-3D.1* were further evaluated in four additional populations with different genetic backgrounds (M3, M2, MS9, and CAW) using the newly designed KASP marker KASP-10 (S5 Table) tightly linked to *QSns.sau-MC-3D.1*. Genotyping was executed for 184, 218, 178, and 388 lines of the M3, M2, MS9, and CAW populations, respectively (S4 Fig).

The M3 population was planted in four different environments, including 2021WJ (M3.F_3_.WJ), 2021CZ (M3.F_3_.CZ), 2022CZ (M3.F_4_.YA), and 2022WJ (M3.F_4_.WJ; Fig 4A). In all the four environments, the group with the homozygous genotype GG from *msf* had significantly greater (*P* < 0.01) SNS than that with the homozygous genotype AA, and the differences between the two groups were 4.13%, 3.59%, 4.90%, and 3.84%, respectively (Fig 4A). The group with the homozygous genotype GG from *msf* had significantly higher (*P* < 0.05) SNS than that with heterozygous genotype GA, and the difference ranged from 2.27 to 3.51% (Fig 4A). Furthermore, there was no significant difference between the group with the homozygous genotype AA and that with the heterozygous genotype GA.

**Fig 4.**
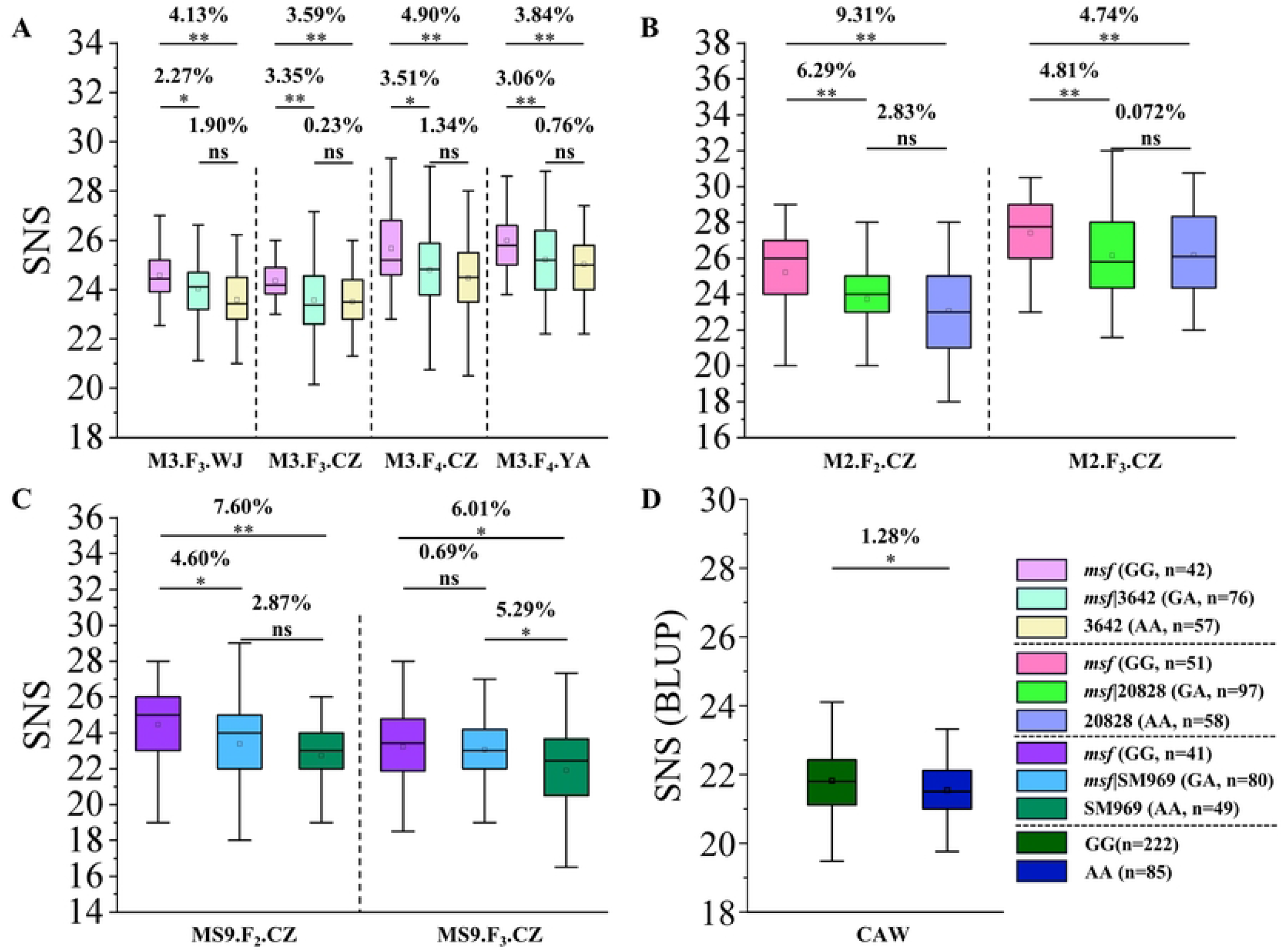
Validation of *QSns.sau-MC-3D.1* in four populations. **A, B, C, and D**, Effects of *QSns.sau-MC-3D.1* in the four validation populations (i.e., *msf* × 3642, *msf* × 20828, *msf* × Shumai969, and CAW). Lines with the homozygous genotype GG of *msf* × 3642, *msf* × 20828, *msf* × Shumai969, and CAW population are 42, 51, 41, and 222, respectively. Lines with the heterozygous genotype GA of *msf* × 3642, *msf* × 20828, and *msf* × Shumai969 population are 76, 97, and 80, respectively. Lines with homozygous genotype AA of *msf* × 3642, *msf* × 20828, *msf* × Shumai969, and CAW population are 57, 58, 49, and 85, respectively. *Significance level at *P* < 0.05, **Significance level at *P* < 0.01, and ns indicates no significant difference between the two groups. Percentage differences between the two groups are indicated above the *P* values at the top of each plot.

The M2 population was planted in two different environments, including 2021CZ (M2.F_2_.CZ) and 2022CZ (M2.F_3_.CZ). In both environments, lines with the homozygous genotype GG from *msf* had significantly higher (*P* < 0.01) SNS than those with AA, and the differences between the two groups were 9.31%, and 4.74%, respectively (Fig 4B). The group with the homozygous genotype GG from *msf* had significantly (*P* < 0.01) greater SNS than that with the heterozygous genotype GA, and the differences between the two groups were 6.29% and 4.81%, respectively (Fig 4B). There was no significant difference between the lines with the homozygous genotype AA and those with the heterozygous genotype GA (Fig 4B).

Likewise, the MS9 population was planted in two different environments, including 2021CZ (MS9.F_2_.CZ) and 2022CZ (MS9.F_3_.CZ). Group 1, with the homozygous genotype GG from *msf*, had a significantly (*P* < 0.01) higher SNS than group 2 (with the homozygous genotype AA) in the two environments with differences ranging between 6.01% and 7.60% (Fig 4C).

In MS9.F_2_.CZ, the group with the homozygous genotype GG from *msf* was significantly (*P* < 0.05) higher SNS than the heterozygous genotype GA group, with a 4.60% significant difference, while in MS9.F_3_.CZ, there was no significant difference (Fig 4C). Moreover, unlike in M3 and M2 populations, there was a significant (*P* < 0.05) difference between the group with the heterozygous genotype GA and the group with the homozygous genotype AA (Fig 4C).

In the CAW population, the group with the homozygous genotype GG from *msf* showed significantly higher SNS than that of the homozygous genotype AA from CN16 (excluding heterozygotes, *P* < 0.05, Fig 4D). The above results indicate that *QSns.sau-MC-3D.1* is a major QTL controlling SNS.

### Effects of *QSns.sau-MC-3D.1* and *WAPO1* on increasing SNS

The effects of *QSns.sau-MC-3D.1* and *WAPO1* on increasing the SNS were further evaluated (Fig 5). Compared with the lines without any of the positive alleles increasing SNS, those only possessing the positive allele GG of *QSns.sau-MC-3D.1* or H2 of *WAPO1* significantly (*P* < 0.01) increased SNS by 2.61% and 3.54%, respectively. And those with the combination of positive alleles of both *QSns.sau-MC-3D.1* and H2 significantly (*P* < 0.01) increased SNS by up to 7.10% (Fig 5). In addition, lines with the combination of positive alleles of *QSns.sau-MC-3D.1* and H2 significantly (*P* < 0.01) increased SNS by 4.37 and 3.44%, respectively, compared to those with either positive allele of *QSns.sau-MC-3D.1* or H2 (Fig 5). However, there was no significant difference between the lines with *QSns.sau-MC-3D.1* and H2 (Fig 5), indicating that the genetic effect between *QSns.sau-MC-3D.1* and *WAPO1* may be additive.

**Fig 5.**
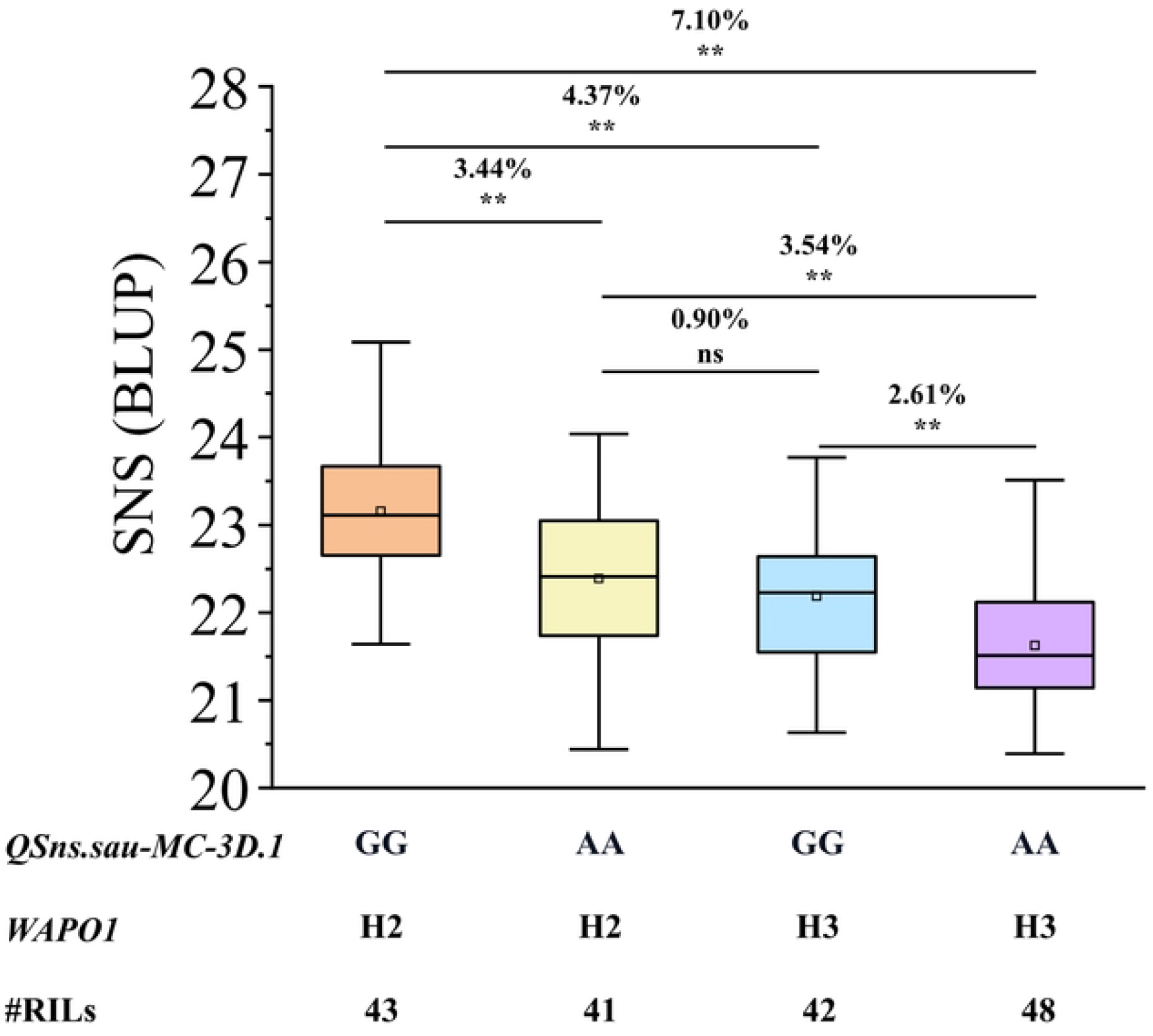
The additive effects of *QSns.sau-MC-3D.1* and *WAPO1* on increasing SNS. ‘H2’ and ‘H3’ represented the H2 (140^T^+115insertion) and H3 (140^G^+115insertion) haplotype of *WAPO1*, respectively. **Significance level at *P* < 0.01, and ns indicates no significant difference between the two groups. Percentage differences between the two groups are indicated above the *P* values at the top of each plot.

### Correlation between major QTL and other agronomic traits

The lines carrying H2 of *WAPO1* in the MC RIL population were removed and the remaining lines were used to detect correlations between *QSns.sau-MC-3D.1* and other yield-related traits. The remaining lines were divided into two groups: lines with the homozygous genotype from *msf* (GG, 42 lines) or CN16 (AA, 48 lines) based on genotyping results using KASP-10 (Fig 6). There were no significant differences (*P* > 0.05) between the two groups for any of the yield-related traits (ETN, PH, SL, AD, and TKW), suggesting that the expression of *QSns.sau-MC-3D.1* was likely independent of these agronomic traits (Fig 6). Similarly, the lines that did not carry the homozygous genotype GG of *QSns.sau-MC-3D.1* were divided into two groups: lines with the H2 (41 lines) or H3 (48 lines) based on genotyping results with K-WAPO1 (Fig 6). There were significant differences between the two groups in ETN and PH (Fig 6), indicating that H2 haplotype of *WAPO1* may affect ETN and PH.

**Fig 6.**
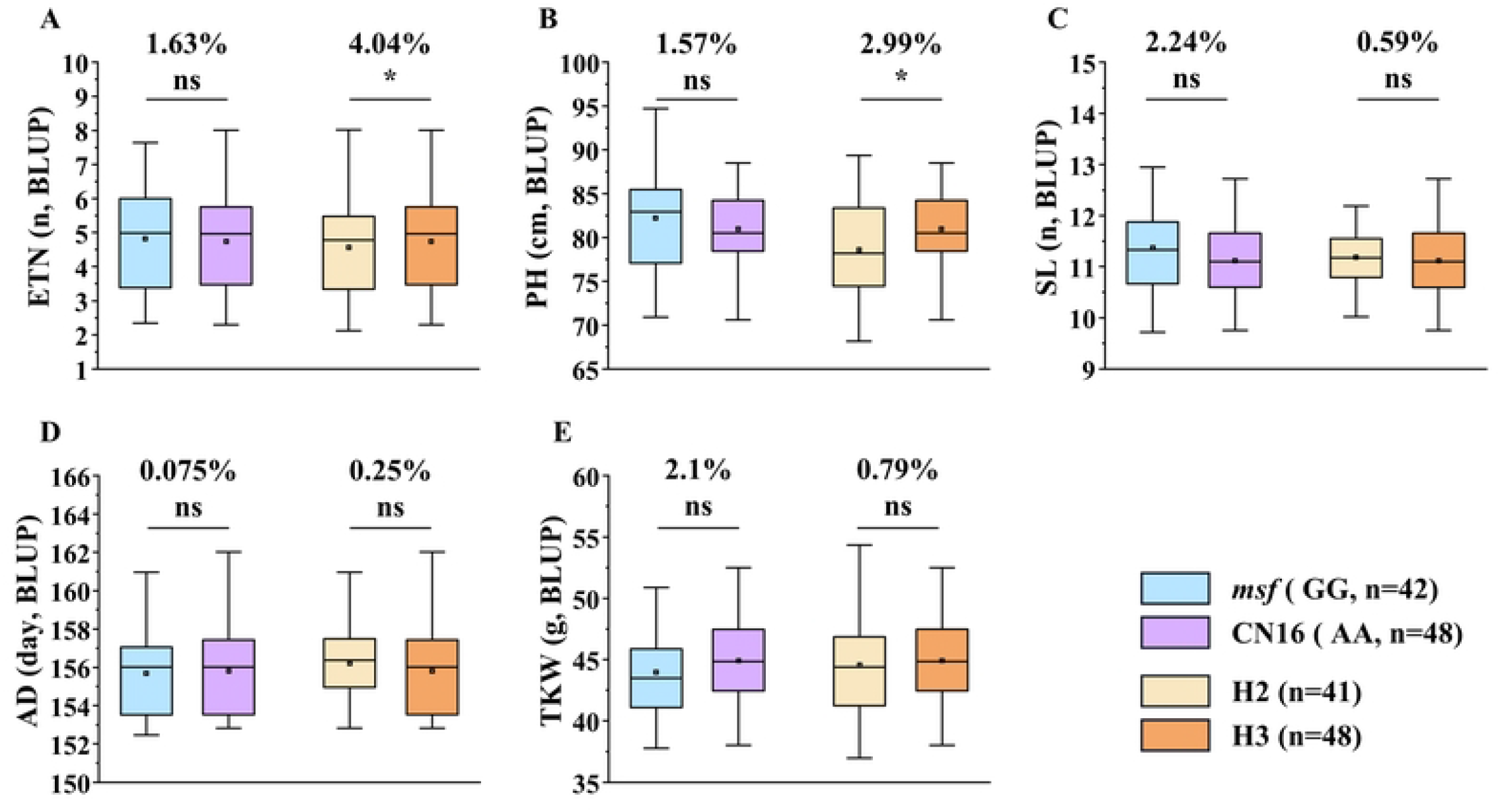
Effects of two major QTL (*QSns.sau-MC-3D.1* and *WAPO1*) on other agronomic traits. **A**, Effective tiller number (ETN); **B**, Plant height (PH); **C**, Spike length (SL); **D**, Anthesis date (AD); **E**, Thousand kernel weight (TKW); *Significance level at *P* < 0.05, ns indicates no significant difference between the two groups. Percentage differences between the two groups are indicated above the *P* values at the top of each plot.

### *QSns.sau-MC-3D.1* underwent positive selection in artificial domestication and breeding

In order to comprehensively and systematically evaluate the distribution of *QSns.sau-MC-3D.1* in Chinese wheat accessions, three hundred and eighty-eight accessions of the CAW population were genotyped using KASP-10. According to the polymorphism of KASP-10, the accessions were divided into two groups in the CAW population: accessions with the homozygous genotype GG and those with AA (excluding heterozygous genotype GA).

In 143 Chinese landraces (ML), the homozygous genotype GG of *QSns.sau-MC-3D.1* was dominant in all seven wheat zones except III (23.53%), V (33.33%), and VII (20%, S5A Fig). A population with 245 Chinese modern cultivars (CMC) was used to further reveal the *QSns.sau-MC-3D.1* distribution in China. As shown in S5B Fig, the frequency of homozygous genotype GG for *QSns.sau-MC-3D.1* was dominant in almost all zones except the V zone (12.5%). I, II, and III belong to the zones with the oldest and strongest wheat breeding programs in China [24]. It’s worth noting that the average frequency of homozygous genotype GG at *QSns.sau-MC-3D.1* in zones I (66.67%), II (86.57%), and III (65%) was 72.74% (S5A Fig), which was much higher than that in ML (62.68%, S5B Fig), suggesting that modern breeding has greatly increased its frequency in CMC. Furthermore, in ML, there was no significant difference in SNS between the group with GG and that with AA (S5C Fig). In CMC, the group with GG had significantly greater (1.58%, *P* < 0.05) values for SNS than that with AA (S5D Fig). This suggest breeders may have indirectly increased the frequency of genotype GG of *QSns.sau-MC-3D.1* in modern breeding by selecting genotypes with higher SNS.

### Identification of candidate gene(s)

There were 93 high-confidence genes within the interval of *QSns.sau-MC-3D.1* (53.61-64.40 Mb, S6 Table). The expression patterns of those genes in various tissues and spikes at different developmental stages were analyzed, and the results showed that there were 9 genes greatly expressed in spike at the reproductive stage and 7 genes highly expressed in spike at the single and double ridge stage with 2 genes shared (S6 Fig), suggesting that these 2 genes might be involved in spike development. *TraesCS3D03G0222600* and *TraesCS3D03G0216800* encoding MYB-like transcription factor and basic helix-loop-helix (HLH) transcription factor, respectively, were likely related to spike development based on gene annotation (S6 Table). qRT-PCR analysis further suggested that only the expression level of *TraesCS3D03G0216800* was significantly enhanced in *msf* (*P* < 0.05, S7 Fig). Taken together, our data suggested that *TraesCS3D03G0216800* may play a regulatory role in determining SNS.

## Discussion

### Phenotypic correlations among investigated traits

In this study, SNS was significantly and positively correlated with SL (Table 3). This is consistent with previous studies [25], suggesting that longer SL provides room for more spikelets to grow [26]. There was a significant positive correlation between SNS and AD (Table 3). This result with previous studies indicated that plants with a longer flowering time may have more time for the differentiation and development of the spikelet primordia [27]. Moreover, SNS was significantly and negatively correlated with TKW (Table 3). Considering the source reservoir relationship in the plant, the increase in the number of spikelets may lead to a decrease in the nutrients allocated to a single kernel [25]. These conclusions provide a vital basis for understanding the complex relationships among wheat yield traits to further improve wheat yield.

### *QSns.sau-MC-3D.1* is a novel QTL for SNS

The physical locations of the QTL/SNP for SNS in previous studies were used for comparing to that of *QSns.sau-MC-3D.1* (S8 Fig). *QSns.sau-MC-3D.1* was located between 53.61 and 64.40 Mb in the deletion bin 3DS6-0.55-1.00 on chromosome arm 3DS in CS (S8A and B Fig), which was different from the previously reported SNS-related QTL/SNP (S8C Fig). For example, *QTsn.cau-3D.3* was located at 3.99 Mb on a chromosome arm 3DS with the peak marker CAP11_c3914_325 [7]. *QTL1935* was physically located on a chromosome arm 3DS at 110.04-129.55 Mb, overlapping with *QSns.cd-3D* [28]. Two SNS-related SNPs, T/C [7] and C/T [29], were located at 512.68 Mb and 600.26 Mb, respectively, on a chromosome arm 2BL. The comparison of the physical locations of *QSns.sau-MC-3D.1* with those of previously reported QTL suggests that *QSns.sau-MC-3D.1* is likely a novel QTL controlling SNS (S8C Fig).

### SNS is not affected by 1BL/1RS translocation

The 1BL/1RS translocated chromosomes have been widely used to develop wheat cultivars [30]. In the current study, CN16 is a cultivar with 1BL/1RS translocation [31]. Identification of 1BL/1RS translocations in the MC RIL population showed 58 lines with 1BL/1RS translocations, while 116 lines with non-1BL/1RS translocations (S9 Fig). *t*-test showed that there was no significant (*P* > 0.05) difference between the SNS of the two groups (S9 Fig), suggesting that translocations of 1BL/1RS chromosomes may not affect SNS in the MC RIL population. However, given its non-recombinant nature and distorted segregation, wheat genotypes used to construct segregating populations should be carefully selected when aiming to identify and clone genes on related chromosomes [31].

### The yield improvement potential of *QSns.sau-MC-3D.1* is likely superior to that of *WAPO1*

Here, two major and stably expressed QTL, *QSns.sau-MC-3D.1* and *QSns.sau-MC-7A* (*WAPO1*) for SNS were identified. Both the positive allele of *QSns.sau-MC-3D.1* and the H2 haplotype of *WAPO1* significantly (*P* < 0.01) increased SNS (Fig 3 and S3 Fig). In previous studies, SNS tends to be significantly and negatively correlated with ETN and TKW [8], and significantly and positively correlated with PH and AD [9], which is not conducive to yield improvement and field breeding. In this study, the expression of *QSns.sau-MC-3D.1* was independent of the above agronomic traits (Fig 6). However, H2 haplotype expression of *WAPO1* may affect ETN and PH (Fig 6).

Furthermore, *QSns.sau-MC-3D.1* underwent positive selection in modern breeding (S5B and D Fig). To sum up, in the process of breeding utilization, the breeding potential of *QSns.sau-MC-3D.1* may be superior to that of *WAPO1*.

### Candidate gene analysis of *QSns.sau-MC-3D.1*

According to the CS reference genome V2.1, there were 93 annotated high-confidence genes within the candidate intervals of *QSns.sau-MC-3D.1* (S6 Table), and spatiotemporal expression patterns and functional annotations of those genes suggest that two genes, *TraesCS3D03G0222600* and *TraesCS3D03G0216800*, may be involved in determining the development of SNS (S6 Fig, S6 Table). Previous studies have also shown that MYB transcription factors determine the fate of spikelet meristem [32,33], and HLH transcription factor regulate flowering time in grasses [34]. However, qRT-PCR of the two genes showed that only *TraesCS3D03G0216800* was significantly expressed between parents. These results suggested that *TraesCS3D03G0216800* may be a candidate gene for *QSns.sau-MC-3D.1* and laid a vital foundation for fine mapping and map-based cloning.

## Materials and Methods

### Plant Materials

A wheat population (MC population) containing 198 F_6_ RILs (excluding two parental lines) was derived from an across between *msf* and Chuannong 16 (CN16) used in this study. *msf* is a spontaneous mutant characterized by multi-spikelets, multi-florets (Fig 7), large spike and high fruiting rate. CN16 is a commercial wheat cultivar, developed by Triticeae Research Institute of Sichuan Agricultural University, with excellent agronomic performances including multiple tillers and good plant type [31]. The MC population was used for QTL identification. Major and novel QTL for SNS identified in the MC RIL population were validated in three populations, including *msf* × 3642 (M3, F_3_, and F_4_, 184 lines), *msf* × 20828 (M2, F_2_, and F_3_, 218 lines), and *msf* × Shumai969 (MS9, F_2_, and F_3_, 178 lines). Line 20828 was kindly provided by Dr. Wu Yu (Chengdu Institute of Biology, Chinese Academy of Sciences). The line 3642 and cultivar Shumai969 were provided by the Triticeae Research Institute of Sichuan Agricultural University. In addition, three hundred and eighty-eight Chinese wheat accessions (CAW), including 143 landraces from the mini-core collection (ML) and 245 modern cultivars (CMC) released since the 1940s (S7 Table) [35], were further employed to verify the effect of the major QTL.

**Fig 7.**
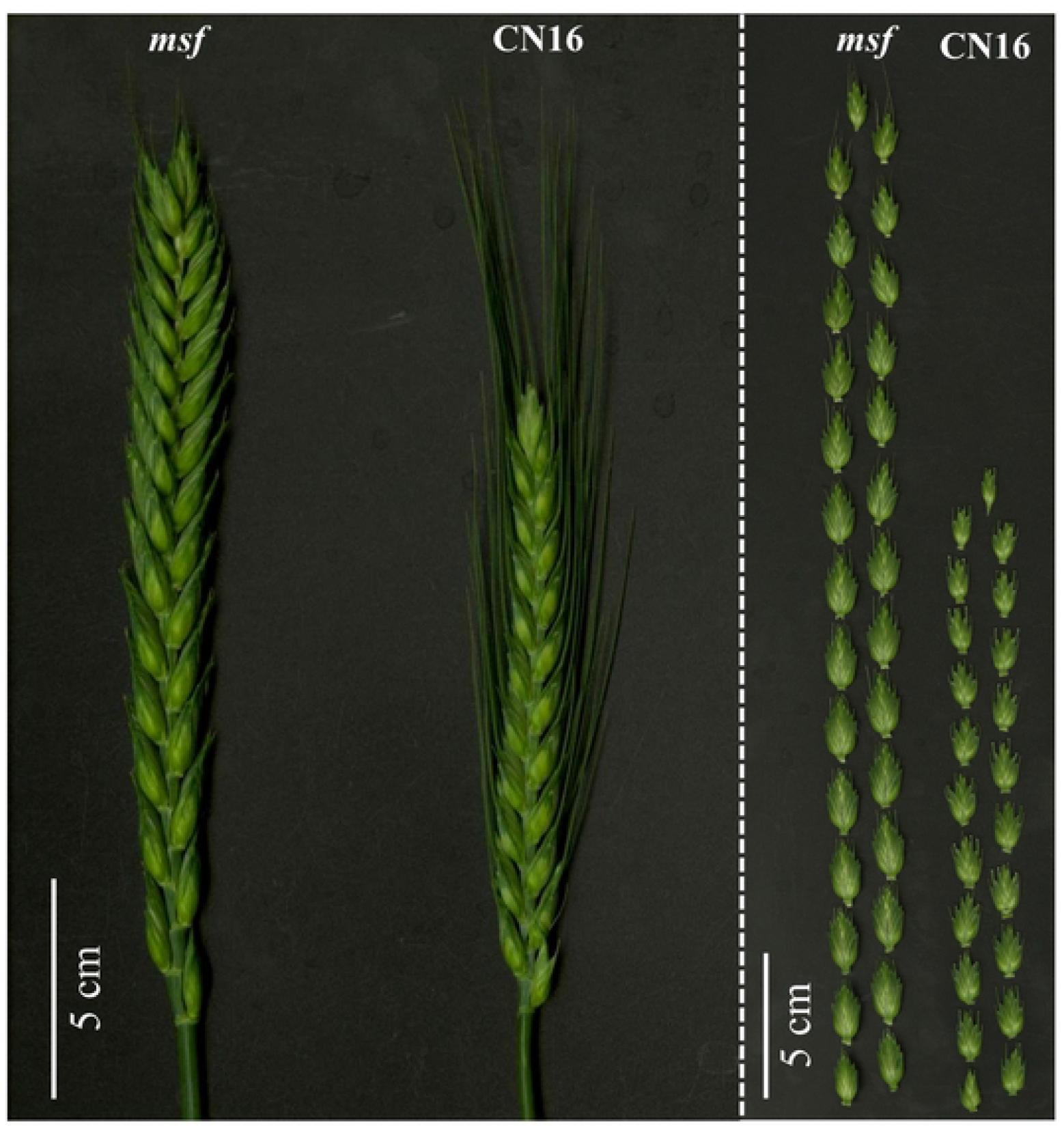
Phenotypes of *msf* and CN16. The white bar represents 5 cm.

### Field experiments and phenotypic evaluation

The MC RIL population and parents were planted in five different environments including Wenjiang (103°51′ E, 30°43′ N) in 2021 and 2022 (2021WJ and 2022WJ); Chongzhou (103°38′E, 30°32′N) in 2021 and 2022 (2021CZ and 2022CZ); Ya’an (103°0′E, 29°58′N) in 2021 (2021YA). The trials in all the environments were performed in a randomized block design with two replications. Seven seeds of each line were planted in a 0.75 m row with 0.1 meters between plants, and 0.3 meters between rows. Field management followed local practices for wheat production.

SNS was measured by counting the number of spikelets of the main spike, effective tiller number (ETN) was counted as the number of the fertile spike per plant before harvest, plant height (PH) was calculated as the distance from the base to the tip of the highest spike (excluding awns) per plant, spike length (SL) was measured as the length from the rachis node of the first base spikelet to the tip of the main spike (excluding awns) per plant, TKW was calculated as 10 times the average weight of 100 kernels in each line, and anthesis date (AD) was defined as the number of days between sowing and 50% of the plants flowering in each line. At least four plants free of disease in each replicate of each line with consistent growth were selected for trait measurement and then averaged for further analysis.

Three segregation populations for validation, M3, M2, and MS9, were planted in four (2021WJ, 2021CZ, 2022CZ, and 2022YA), two (2021CZ and 2022CZ), and two (2021CZ and 2022CZ) different environments, respectively. CAW was planted in three different environments including Luoyang (Henan province, China) in 2002 and 2005 (2002 LY and 2002 LY), Shunyi (Beijing, China) in 2010 (2010 SY), and Chongzhou (103°38′E, 30°32′N) in 2022 (2022CZ). Planting trials and phenotypic traits collection of CAW (2002 LY, 2005 LY, and 2010 SY) were described by Wang et al. [35] and Zheng et al. [24], respectively. The methods of planting and SNS measurement for M3, M2, MS9, and CAW (2022CZ) were same as for the MC RIL population.

### Genotyping

Genomic DNA extraction from leaf samples collected at the joining stage adopted the CTAB protocol [36], and DNA quality was assessed using a NanoDrop One C (Thermo Fisher Scientific, Assembled in the USA). The 198 lines and parents of the MC population were genotyped using the Wheat 16K SNP array from Mol Breeding Company (Shijiazhuang in Hebei province; http://www.molbreeding.com). The Wheat 660K SNP array from Capitalbio Technology (Beijing, http://www.capitalbiotech.com/) was also used to genotype the two parents of the MC RIL population. The primers used in this study were synthesized by Tsingke Biotechnology Co., Ltd. (https://www.tsingke.com.cn/).

### Data analysis

The frequency distribution of SNS in each environment and correlation analysis were performed using Origin 9.0 software and SPSS V26.0 for Windows (SPSS Inc., Chicago, IL), respectively. The best linear unbiased prediction (BLUP) dataset for all the investigated traits was tested using SAS V8.0 (SAS Institute, Cary, North Carolina). The calculation of the broad-sense heritability (*H*^2^) of SNS was performed as described by Smith et al. [37]. Analysis of variance (ANOVA) was performed using the Aov (ANOVA of multi-environment trials) module of QTL IciMapping V4.1 (https://www.isbreeding.net/) to detect interactions between replications, genotypes and environments. The student’s *t*-test performed by SPSS V26.0 was used to evaluate the differences in parents and RIL population. Furthermore, the correlation coefficients between traits were calculated using SPSS V26.0 based on the BLUP dataset of each trait.

### Linkage map construction and QTL analysis

14,870 SNP markers (14,868 mSNP segments + 2 polymorphic SNP) from the 16K SNP array and the 660K SNP array were obtained. Firstly, the minor allele frequency (MAF) was calculated for each SNP marker in the MC RIL population, and those with MAF greater than 0.3 were retained. Secondly, the retaining markers were analyzed by using the BIN function in QTL IciMapping V4.1, based on their segregation patterns in the MC RIL population, with parameters ‘distortion value’ and ‘missing rate’ being set as 0.01 and 20%, respectively. A single marker with the lowest ‘missing rate’ from each set of bin markers was further selected. Finally, the bin markers were grouped and sorted using the Kosambi mapping function in QTL IciMapping V4.1 with the logarithm of odds (LOD) greater than 3 after preliminary analysis of markers with LOD scores ranging from 2 to 10. The finally retained markers were used to generate genetic maps using the ‘MAP’ function in the QTL IciMapping V4.1 software and maps were further drawn in MapChart V2.32. The flanking sequences (200bp) of SNPs were used to blast against (*E*-value of 1e^-5^) genome sequences of the International Wheat Genome Sequencing Consortium (IWGSC) Chinese Spring (CS) RefSeq V2.1 [38] to get their physical locations. The syntenic relationships between the genetic and physical maps of the bin markers were presented using the Strawberry Perl V5.24.0.1.

Inclusive composite interval mapping with the biparental population module (mapping method: ICIM-ADD. Step = 1 cM, PIN = 0.001, and LOD threshold = 2.5) in QTL IciMapping V4.1 was performed to detect QTL for SNS.

Among the QTL detected in more than three environments (including BLUP dataset) and explaining greater than 10% of the PVE were considered as major and stable QTL, and those with common flanking markers were treated as identical ones. The detected QTL were basically named as per the International Rules of Genetic Nomenclature (http://wheat.pw.usda.gov/ggpages/wgc/98/Intro.htm). ‘Q’, ‘SNS’, ‘sau’, and ‘MC’ represent ‘QTL’, ‘Spikelet Number per Spike’, ‘Sichuan Agricultural University’, and ‘the MC RIL population’, respectively.

### Comparison with previously reported QTL/SNP for SNS

Previously reported closely linked marker sequences of QTL/SNP related to SNS were obtained from WheatQTLdb V2.0 [6], and further blasted against genomes sequences of IWGSC RefSeq V2.1 [38] to get their physical locations.

### Marker development and QTL validation

To further narrow down the intervals of major QTL, significant SNPs from the 660K SNP array were converted into KASP markers (S5 Table) to genotype the MC RIL population. According to QTL mapping results, the flanking markers closely linked to novel and major QTL were converted to KASP markers as previously described [31]. The validation populations, M3, M2, MS9, and CAW, were genotyped using the KASP marker (S5 Table). The 10 μl reaction system includes 1 μl DNA, 2.6 μl RNA-free deionized water, 5 μl SsoFast EvaGreen mixture (Bio-Rad, Hercules, CA, USA), and 1.4 μl of mixture forward and reverse primers. All KASP processes were carried out on a CFX96 Real-Time PCR Detection System (BioRad, USA) [39]. The lines were divided into three groups based on the genotyping results: (1) lines with homozygous genotype GG from *msf*; (2) lines with homozygous genotype AA from alternative parent; (3) lines with the heterozygous genotype GA. Finally, we assessed the differences in SNS between the three groups using an independent samples *t*-test (*P* < 0.05) to determine the effects of major QTL.

### Identification of lines carrying 1BL/1RS translocation

The parental CN16 is a genotype carrying the 1BL/1RS translocation [31]. Thus, we identified 1BL/1RS translocations of the RILs derived from *msf* and CN16. Firstly, SNP markers on chromosome 1B were screened from the 16K SNP array in the MC RIL population (2,061 markers in total). The markers mapped on 1BS of IWGSC RefSeq V2.1 [38] were identified (501 markers). Secondly, SNP markers genotyped as ‘NA’ (no genotype detected) in CN16 were retained (276 markers) for further analysis. The ‘NA’ information present under 276 markers for each line was counted. According to the distribution of NA in each line, the lines with less than or equal to 19 NA in these 276 markers were considered as non-1BL/1RS translocation lines and those with the number of NA greater than or equal to 62 were 1BL/1RS translocation lines. Moreover, to validate the 1BL/1RS translocation in the MC RIL population, we also used the 1BS- and 1RS-specific markers to detect the translocation [40]. The 20 μl reaction system included 2 μl DNA, 6 μl RNA-free deionized water, 10 μl 2×Taq PCR PreMix (+Blue dye, innovagene), and 1 μl of each primer (10μm). The reaction conditions were as follows: pre-denaturation at 95 °C for 5 min; a total of 35 cycles of denaturation at 95 °C for 30 s, annealing at 62 °C for 30 s and extension at 72 °C for 30 s; and final extension at 72 °C for 7 min. Primer information was listed in S5 Table. Finally, the lines carrying 1BL/1RS translocation from the MC RIL population were counted based on the above two methods.

### Potential candidate gene(s) for major QTL

According to the mapping result, the sequences of the flanking markers were used to blast (*E*-value of 1e^-5^) against the IWGSC RefSeq V2.1 to obtain their physical locations. The high-confidence genes within the physical positions were obtained from WheatOmics 1.0 (http://202.194.139.32/) [41]. The functional annotations of predicted genes were assigned based on UniProt (http://www.uniprot.org/). Gene expression data in various tissues was extracted from expVIP (http://www.wheat-expression.com/). The data on gene expression patterns in different stages of spike development were obtained from a previous study [42]. Furthermore, the expression pattern of the predicted gene was represented in the HeatMap drawn on Hiplot [43].

### Gene expression studies

Total RNA extracted from freshly harvested spikes at single ridge end-stage with the RNAprep pure Plant Kit (Biofit Biotechnologies co. Ltd, Chengdu, China) was digested with RNase-free DNase (Takara) to remove residual genomic DNA. The RNA was reverse-transcribed into cDNA by using a Prime ScriptTM RT Reagent Kit (TaKaRa, Kyoto, Japan) according to the manufacturer’s instructions. SYBR qPCR Master Mix kit (Q711, Vazyme, Nanjing, China) and a Bio-Rad CFX96 real-time PCR detection system (Bio-Rad, Hercules, USA) were used for qRT-PCR. Three biological replicates were performed for each parent, and each sample was assayed three times. The PCR reaction mixture contained: 2 μl cDNA, 5 μl 2X SYBR Green mix, 0.5 μl forward primer, 0.5 μl reverse primer and 2 μl ddH_2_O, in a final volume of 10 μl. The PCR program was as follows: 94 °C for 5 min, followed by 35 cycles of 94 °C for 30 s, 62 °C for 30 s, and finally 72 °C for 30 s. The 2^-ΔΔCt^ method was used to calculate the relative expression levels of the candidate genes. The Actin gene was used as an internal control. Specific primers for qRT-PCR were designed in NCBI and the details of primers were listed in S5 Table.

## Acknowledgments

We thank the anonymous referees for critical reading and revising this manuscript.

## Author Contributions

JGZ finished the study and wrote this manuscript. WL participated in field work and analyzed data. YYY, XLX, JJL, and MD helped phenotype measurement and data analysis. YLL, HPT, QX and QTJ did field work and data analysis. GYC, PFQ, YFJ, and GDC collected and analyzed data. YJH, YR, LWT, and LLG helped with data analysis. YLZ revised the manuscript. YMW discussed results and revised the manuscript. JM designed the experiments, guided the entire study, participated in data analysis, wrote, and extensively revised this manuscript. All authors participated in the research and approved the final manuscript.

## Supporting information

**S1 Table. Sequence information of 5991 SNP markers**.

(XLSX)

**S2 Table. The blastn results of 5991 SNP markers sequences against the reference genome sequence IWGSC RefSeq V2.1**.

(XLSX)

**S3 Table. Comparison of the genetic and physical positions of the bin markers**.

(XLSX)

**S4 Table. Analysis of variance for spikelet number per spike (SNS) in the *msf* × CN16 population**.

(XLSX)

**S5 Table. Details of primers used in this study**.

(XLSX)

**S6 Table. Predicated genes in the interval of *QSns.sau-MC-3D.1***.

(XLSX)

**S7 Table. The information of three hundred and eighty-eight Chinese wheat accessions (CAW)**.

(XLSX)

**S1 Fig. Phenotypic distribution of spikelet number per spike (SNS) at five environments and BLUP**.

(DOCX)

**S2 Fig. Haplotype identification of *WAPO1* in *msf* and CN16**.

(DOCX)

**S3 Fig. Genetic map of the major QTL *QSns.sau-MC-7A* and the effect of *WAPO1***.

(DOCX)

**S4 Fig. Fluorescence PCR genotyping results of the KASP marker KASP-10 in four populations**.

(DOCX)

**S5 Fig. Distribution of 143 Chinese landraces (A) and 245 modern cultivars (B) in ten production zones**.

(DOCX)

**S6 Fig. Expression pattern of genes within the *QSns.sau-MC-3D.1* interval**.

(DOCX)

**S7 Fig. Expression of *TraesCS3D03G0222600* and *TraesCS3D03G0216800* in the spike of parent msf and CN16**.

(DOCX)

**S8 Fig. *QSns.sau-MC-3D.1* comparison with previously reported spikelet number per spike (SNS)-related quantitative trait loci (QTL) and single nucleotide polymorphisms (SNPs)**.

(DOCX)

**S9 Fig. The effect of 1BL/1RS translocations on spikelet number per spike (SNS) in the *msf* × CN16 population**.

(DOCX)

**S1 Data. Data A**. Supporting data for Fig 1 and Fig 2. **Data B**. Supporting data for Fig 3B and S3B Fig. **Data C**. Supporting data for Fig 4A-4C. **Data D**. Supporting data for Fig 5 and Fig 6. **Data E**. Supporting data for S5A-5D Fig. **Data F**. Supporting data for S6 Fig and S9 Fig. **Data G**. Supporting data for Table 3 and S7 Fig.

(XLSX)

## Data Availability Statement

All data are presented in the text and supplementary materials. The raw data for all figures and Supplemental Tables are available in S1 Data file.

